# Self-limiting population genetic control with sex-linked genome editors

**DOI:** 10.1101/236489

**Authors:** Austin Burt, Anne Deredec

## Abstract

In male heterogametic species the Y chromosome is transmitted solely from fathers to sons, and is selected for based only on its impacts on male fitness. This fact can be exploited to develop efficient pest control strategies that use Y-linked editors to disrupt the fitness of female descendants. In simple “strategic” population models we show that Y-linked editors can be substantially more efficient than other self-limiting strategies and, while not as efficient as gene drive approaches, are expected to have less impact on non-target populations with which there is some gene flow. Efficiency can be further augmented by simultaneously releasing an autosomal X-shredder construct, in either the same or different males. Y-linked editors may be attractive option to consider when efficient control of a species is desired in some locales but not others.

## Introduction

The most widely used genetic approach to pest control thus far has been the mass release of sterile males (Alphey et al. 2010). This approach requires inundating the target population with males that have been sterilized by radiation, infected with *Wolbachia*, or modified by transgenesis (Alphey 2014), and has been used successfully against some agricultural pests and disease vectors. However, the numbers released typically have to be at least 10-fold larger than the target population, and sustained for multiple generations, and the associated costs of mass production limit the range of species for which this approach is suitable.

In principle, the introduction into populations of genetic constructs showing preferential inheritance (“gene drive”) may be substantially more efficient than inundative approaches like the sterile insect technique, requiring only small inoculative releases to suppress a population (e.g., less than 1% of the target population) (Burt 2014, Godfray et al. 2017), and promising proof-of-principle demonstrations of such constructs have been reported for malaria-transmitting mosquitoes (Galizi et al. 2014, Galizi et al. 2016, Hammond et al. 2016). One reason for the predicted efficiency is that natural processes of dispersal and migration can be exploited to introduce the construct (and its suppressive effect) from the population into which it was released into other populations that may be more difficult to access (Beaghton et al. 2016, Eckhoff et al. 2017).

In some cases it may be desirable to control a pest species in one location but not in another – for example, to control an agricultural pest in a farmer’s field but not in a nature reserve, or to control an invasive species where it is invasive but not in its native range. In these cases, depending on the biology of the species involved, one may want to consider interventions that are more efficient than mass release of sterile males, but will not have an appreciable impact on non-target populations, even if there is some gene flow. Possibilities include releasing males that carry constructs that kill female descendants (Fu et al. 2010) or cause them to have predominantly male offspring (Galizi et al. 2014), both of which can, in some circumstances, be more efficient than release of sterile males and are still expected to have geographically restricted impacts. If the goal is to modify the target population rather than suppress it, then there is a range of options, some of which should allow geographical targeting (Marshall and Hay 2012, Dhole et al. 2017).

In this paper we explore the possibility of using a construct inserted on the Y chromosome to make edits in autosomal or X-linked genes needed for female survival or reproduction. The key idea is that the construct acts in males to reduce the fitness of female descendants, but since the Y is not found in those descendants, the construct will not be selected against by the harm it causes. This idea was previously suggested by Deredec et al. (Deredec et al. 2011) as a way to increase the efficacy of a driving Y chromosome, but clearly it can also be used in a non-driving context. We use relatively simple “strategic” models to explore some of the features of this approach, starting with an idealized case and comparing its efficiency to that of other self-limiting constructs. We then examine the impacts of various deviations from the ideal, to assess the robustness of the approach, and the impacts of gene flow both into and out of the target population. Finally, we explore how efficiencies can be further increased by including another transgene in the releases to boost the frequency of the modified Y, and discuss some of the molecular options for building these constructs. For ease of exposition, we emphasize results from a set of exemplar scenarios that we consider most illuminating, rather than an exhaustive analysis of a simple model that would need to be made more realistic for any particular use case. Our modeling demonstrates that Y-linked genome editors (YLEs) have a unique combination of features and should be a useful addition to the menu of options to be considered in designing and developing pest control programs.

## Results

### Ideal case and comparison with alternatives

We first consider the release of males carrying an idealized construct in which the only effect of the YLE is to induce 100% knock-out of a target gene, which may be either autosomal or X-linked, and the only effect of the knock-out is complete dominant female lethality. For comparison, we also model idealized versions of several other strategies, including release of males homozygous for:

1. A dominant autosomal lethal gene; this is the bisexual RIDL (“bi-RIDL”) approach and, at least in our model, is equivalent to the release of radiation-sterilized males or of males carrying a *Wolbachia* strain that renders them incompatible with the target population (Carvalho et al. 2015, Mains et al. 2016, Zhang et al. 2016).
2. A dominant autosomal female-specific lethal gene (“fs-RIDL”); males inheriting the gene have normal fitness and transmit it to the Mendelian 50% of progeny (Schliekelman and Gould 2000, Thomas et al. 2000). In some species it might be possible to use repressible autosomal copies of a male-determining gene for this strategy (Criscione et al. 2016, Krzywinska et al. 2016).
3. A dominant autosomal female-specific lethal showing drive in males (“fs-RIDL-drive”); males inheriting the gene have normal fitness and transmit it to 100% of progeny (Thomas et al. 2000, Hammond et al. 2016). One possible implementation involves linking a male-determining gene to a male-limited gene drive system (Adelman and Tu 2016, Piaggio et al. 2017).
4. A dominant autosomal gene causing all sperm to carry the Y chromosome, and therefore all progeny to be male (“X-shredder”) (Schliekelman et al. 2005, Galizi et al. 2014, Galizi et al. 2016).

The efficiency of these alternative control strategies was compared using a simple deterministic model of a population with discrete non-overlapping generations, random mating, male heterogamety, separate juvenile and adult stages, and density-dependent mortality occurring in the juvenile stage only. For the YLEs and alternatives (1)-(3), we considered variants where the lethal gene acts before density-dependent mortality (e.g., at the embryonic stage), or after (e.g., at the juvenile-adult transition), and for all of these except (1) we also considered the case where the gene causes female sterility rather than death, as such differences have previously been shown to impact the efficiency of control (Phuc et al. 2007, Yakob et al. 2008, Deredec et al. 2011, Alphey and Bonsall 2014). In all cases we assumed that only adult males are released, that they have survival and mating success equal to the wildtype males, and that a constant number is released each generation (rather than, for example, a constant proportion of a declining target population).

Figure 1 shows the population size (number of females) over time with recurrent releases of the different alternative constructs into a target population with an intrinsic rate of increase of *R_m_*=6 and release rates of 10% or 50% of the initial male population size. In both cases the different types of YLE eliminated the population, with no difference whether the target gene was autosomal or X-linked, with the most rapid decline occurring if the edit caused death after density-dependence (hereafter denoted a YLE-a construct). Constructs that combined dominant female lethality and male drive were equally good as the YLEs (indeed, indistinguishable), and all other approaches were less effective.

**Fig. 1.**
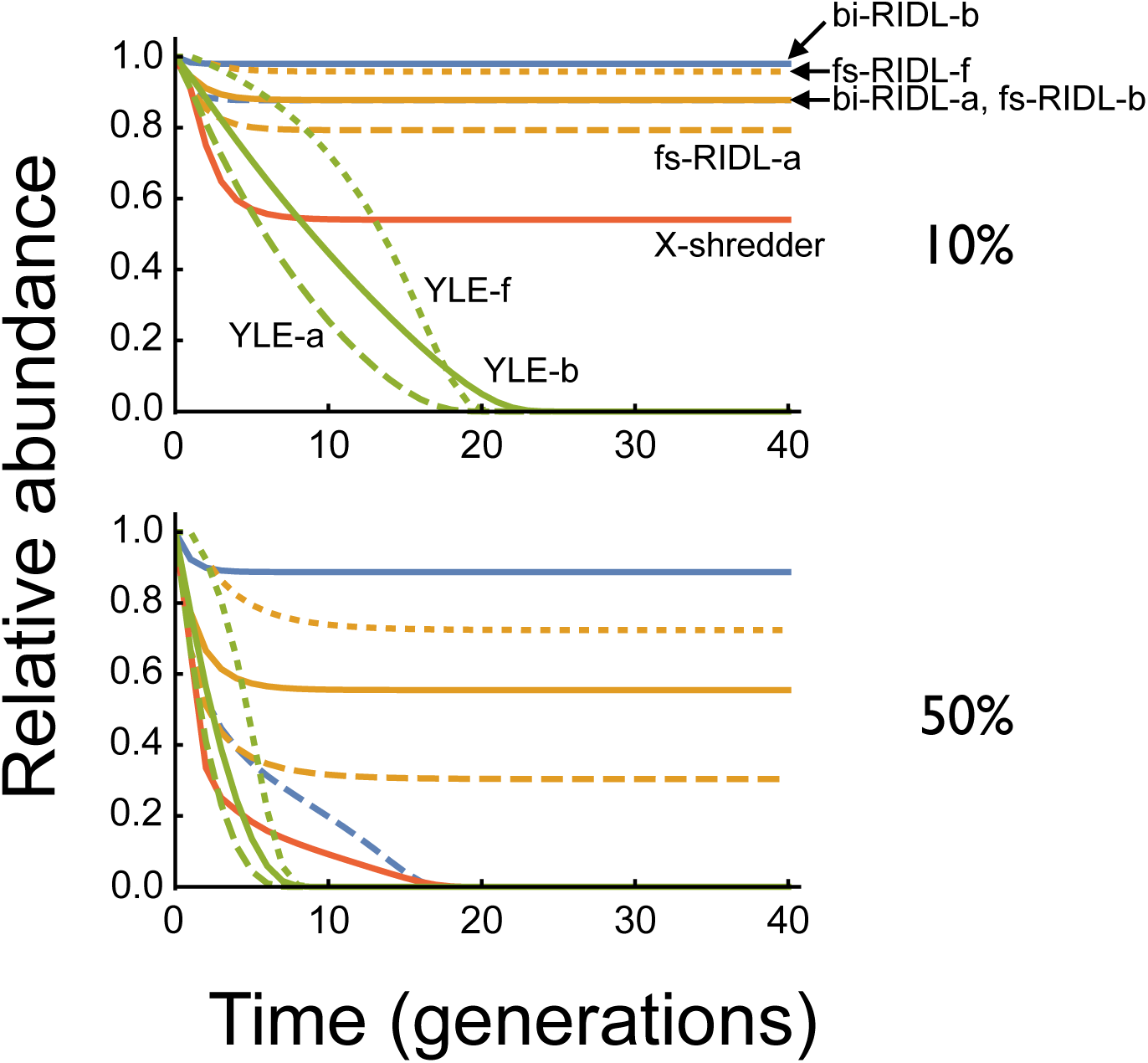
The time course of population control with different self-limiting constructs. The number of females in the population, relative to the pre-release number, is plotted against the number of generations of releases. In each generation an equal number of males is released, either 10% of the initial male population (top), or 50% (bottom). Alternative male genotypes include bi-RIDL (blue), fs-RIDL (orange), X-shredder (red), and YLE (green), and alternative modes of action include death before density-dependent mortality (suffix -b, solid lines), death after density-dependence (-a, dashed), or female sterility (-f, dotted). The three fs-RIDL-with-drive strategies were indistinguishable from the three YLE strategies. *R_m_*=6.

One measure of the efficiency of a construct is the release rate required to achieve a specified level of control in a specified timeframe. Figure 2 shows the release rates required to suppress the number of females in a population by 95% as a function of the duration of the intervention program for the different alternative constructs and different values of *R_m_*. As expected, the required release rates decline as the allowed number of generations increases, and larger releases are needed when *R_m_* is higher. Again, YLEs and female lethality, male drive constructs were indistinguishable, and consistently more efficient than the alternatives, with the differences increasing with both the duration of control and *R_m_*. For a YLE-a construct, the required release rates for different levels of suppression and different values of *R_m_* are shown in Table 1, assuming 36 generations of releases. In the easiest scenario considered (67% suppression with *R_m_*=2), release rates of only 1% of the initial population are needed, whereas for the most difficult scenario (99% suppression with *R_m_*=12), release rates of 5.8% are needed. Sterile male releases would need to be one or two orders of magnitude larger (Table 1).

**Fig. 2.**
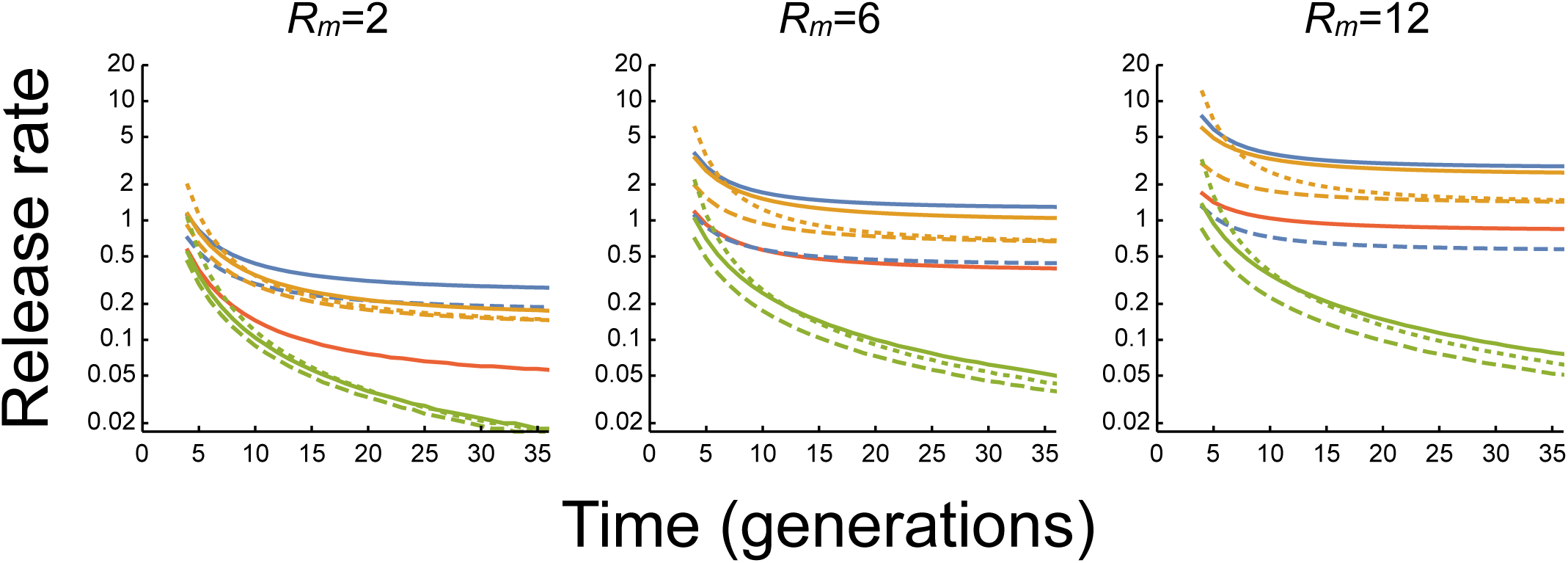
Release rates required to suppress the number of females in a target population to 5% of its initial value as a function of the numbers of generations of releases. Each line represents an alternative strategy; colour and style of lines as in Fig. 1. Release rates calculated as a proportion of the initial number of males; note log scale.

**Table 1.** Release rates required to control a population with a Y-linked editor as a function of the desired level suppression and the intrinsic rate of increase of the population (*R_m_*). Release rates are for the idealized case with the target gene causing female-specific lethality after density dependence, and assuming releases occur for 36 generations. Numbers in parentheses indicate how many times larger the release rate would have to be to achieve the same level of control with sterile males (bi-RIDL-b).

| Level of<br>suppression | $R_m$ | | |
| --- | --- | --- | --- |
|  | 2 | 6 | 12 |
| 67% | 0.010 (27) | 0.021 (58) | 0.026 (108) |
| 95% | 0.015 (18) | 0.036 (35) | 0.050 (56) |
| 99% | 0.016 (16) | 0.041 (32) | 0.058 (49) |

#### Sensitivity to deviations from ideal case

The idealized parameter values we have considered thus far may be difficult to achieve in practice, and therefore it is important to consider how efficiency will be affected by less than perfect performance. As an example, figure 3 shows how the release rates required for 95% suppression in 36 generations are affected by changes in six molecular and fitness parameters, leaving the others at their baseline values. Release rates are not much affected by decreasing the knock-out rate, as long as it remains above 0.9, and not much affected by decreasing the dominance coefficient of female lethality, or the homozygous effect on female survival, as long as they remain above 0.8. If the parameters fall much below these thresholds, then population control may fail, particularly for higher *R_m_*. It makes little difference in these cases whether the target gene is X-linked or autosomal. By contrast, if the knock out also affects male fitness, then the required release rates increase if the target gene is autosomal but not if it is X-linked. Males with the YLE transmit X-linked targets only to their daughters, and if those die, the edit does not appear in male descendants, so male fitness effects do not matter. Finally, if the YLE itself reduces male fitness, then release rates will need to increase.

**Fig. 3.**
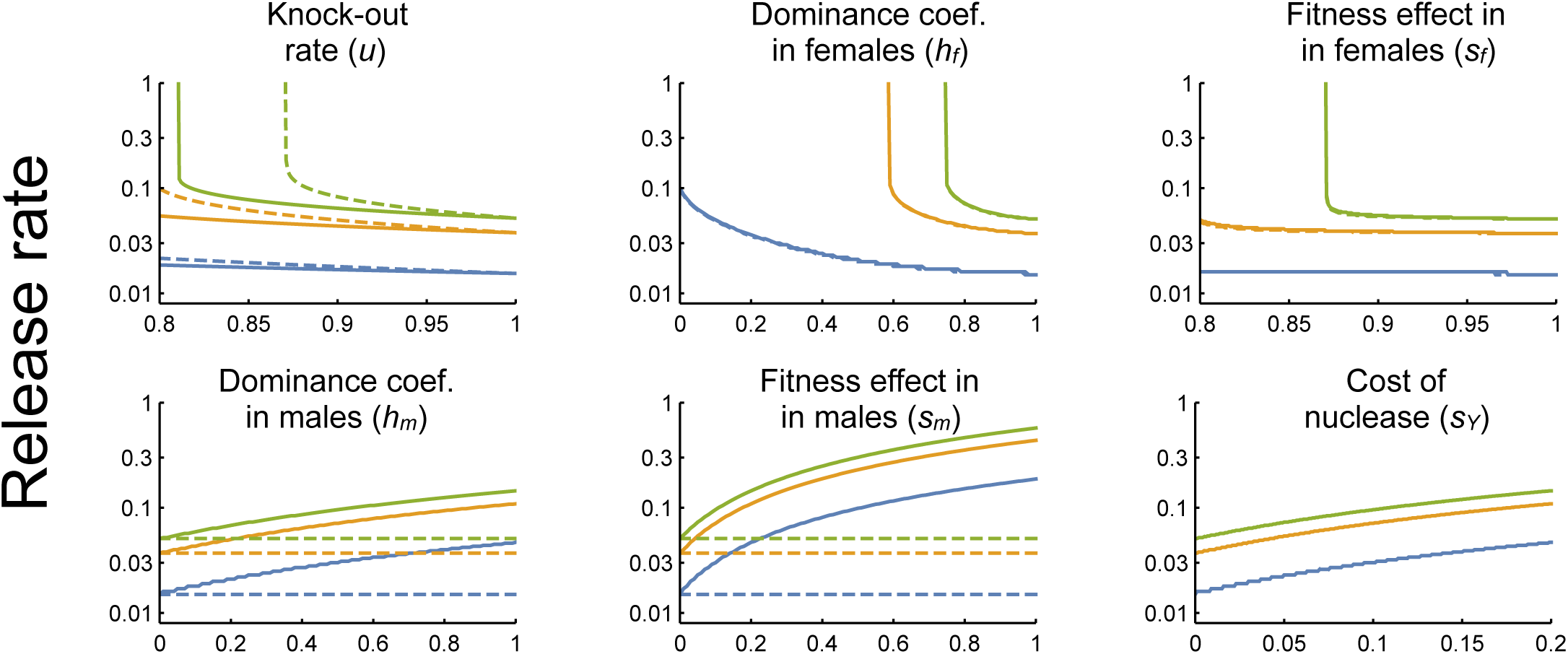
Release rates required to suppress the number of females in a target population to 5% of its initial value in 36 generations with a YLE-a construct as a function of various genetic and fitness parameters, for *R_m_* = 2, 6, and 12 (blue, orange and green, respectively). Solid lines are for an autosomal target locus, and dashed lines for X-linked (when not visible, they are indistinguishable from the solid lines). When varying *h_m_*, *S_m_* was set to 0.2, and when varying *S_m_*, *h_m_* was set to 1; otherwise, all parameters set to baseline ‘idealized’ values. Release rates higher than those shown always give greater suppression after 36 generations, except when the target is an autosomal locus and the editing rate (*u*) less than 1 — in this case higher release rates can give an initially faster decline, but a higher equilibrium number of females.

#### Effect of halting releases

Because selection against YLEs is absent or weak, the consequences of stopping releases before population elimination are markedly different than with the other self-limiting strategies, where the population rapidly recovers (Fig. 4a). In the idealized case of a YLE with no effect on male fitness, halting releases stops the population decline, but it does not recover, and the population remains suppressed indefinitely (Fig. 4b). With a YLE-a the equilibrium abundance of females in our model is:

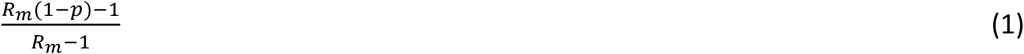

where *p* is the proportion of males with the YLE. Note that if *p* > 1-1/*R_m_* the population is eventually eliminated without further releases. Alternatively, if the YLE imposes a small fitness cost on males (*s_Y_* > 0), then it will be slowly lost after releases are stopped, and the population slowly recover (Fig. 4c). If the release rate (*η*) is above a threshold value

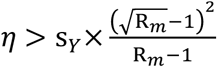

then the population is eventually eliminated, whereas for release rates below this threshold the equilibrium frequency of a YLE-a is:

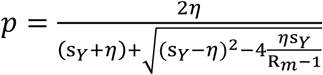

with the equilibrium abundance of females given by equation (1).

**Fig. 4.**
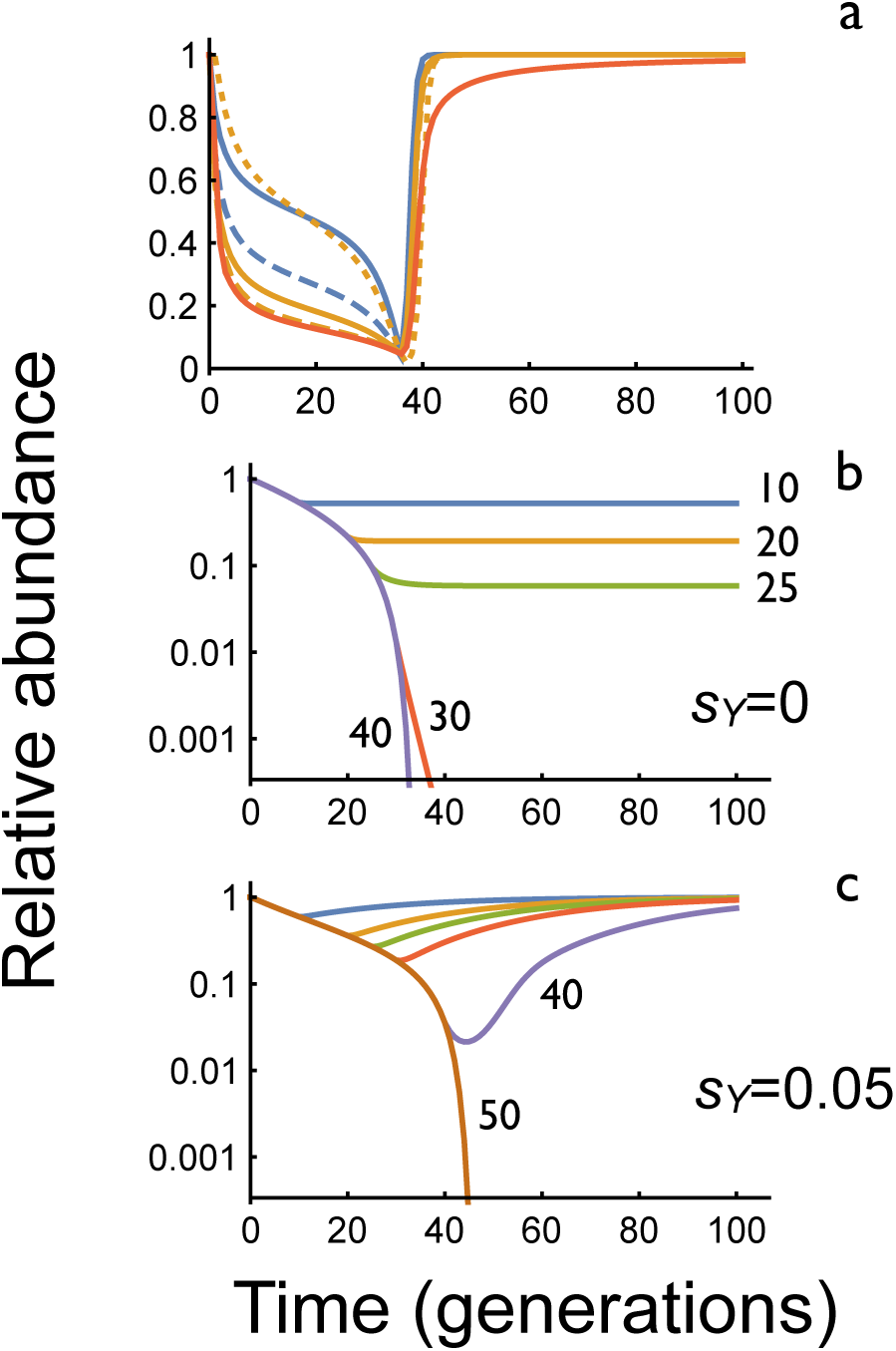
The effect of halting releases. a. Relative abundance of females after 36 generations of releases of RIDL, fs-RIDL, or X shredder constructs. Colour scheme same as Fig. 1. For each construct the release rates were chosen to give 95% suppression after 36 generations. In all cases populations recover rapidly after releases stop. b, c. Relative abundance of females after releases of YLE-a constructs with releases stopping after the indicated number of generations with *s_Y_*=0 (b) or *s_Y_*=0.05 (c). Release rates are 0.05 with an autosomal target gene; note logarithmic Y-axis.

#### Incorporating spatial structure with gene flow

We have thus far considered a single closed, random mating population. We now consider two populations, one targeted for control and the other not, with some migration between them, and ask two questions: What effect does immigration of wildtype organisms into the target population have on the level of control, and what effect does control of the target population have on the non-target population?

To investigate the impact of immigration into the target population, we assume both males and females immigrate, and this occurs at the adult stage, after density-dependent mortality. Females may mate either before or after immigrating. We initially assume movement is unidirectional, so all immigrants are wildtype (also equivalent to assuming the non-target population is very large). The effect of immigration on the release rates required to achieve 95% suppression in 36 generations with a YLE-a are shown in Figure 5. Obviously, if the number of immigrants each generation is equal to a proportion *g* of the initial population size, then no matter how many males of whatever genotype are released, it will not be possible to reduce the population below *g* (unless immigration rates or the source population are also controlled). And immigration of mated females presents more of a problem than immigration of unmated females (Prout 1978).

**Fig. 5.**
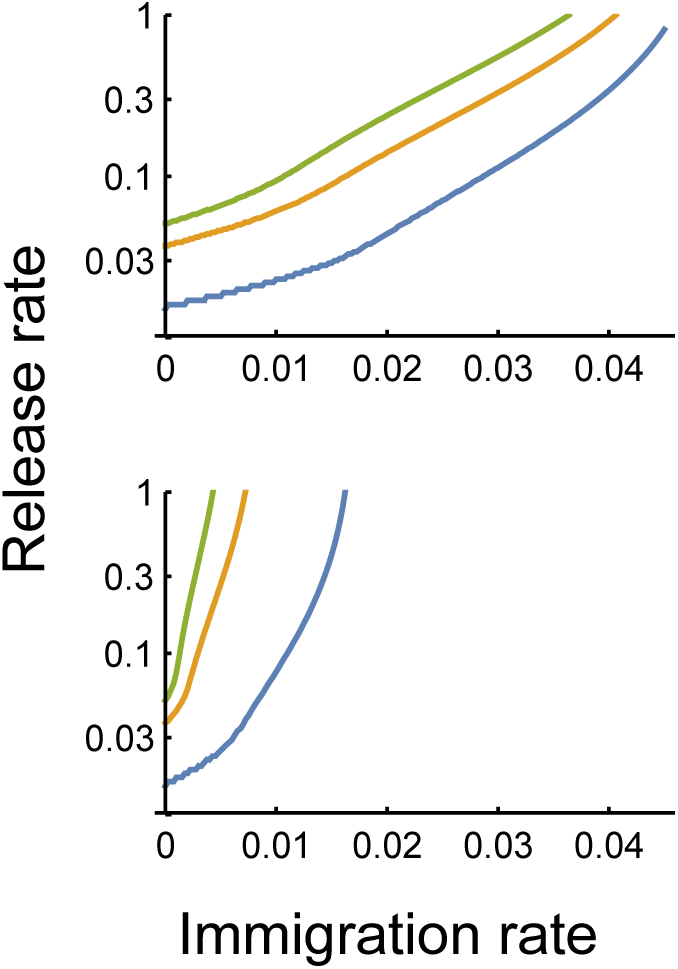
Release rates required to suppress the number of females in a target population to 5% of its initial value in 36 generations with a YLE-a construct as a function of the immigration rate, for *R_m_* = 2, 6, and 12 (blue, orange and green, respectively). Top: immigration of males and unmated females. Bottom: immigration of males and mated females. Results are the same whether target locus is autosomal or X-linked.

To analyze the impact of gene flow from the target to non-target populations, consider first the numbers of transgenics in the target population. For example, a release rate of 3.7% of an ideal YLE-a for 36 generations will suppress a population by 95% (Table 1). The total number of YLE males in the target population over this time is about 13 (i.e., 13 times the original number of males), with a maximum number in any one generation of about 0.5 (Fig. 6). If, for example, the probability that any one of these males emigrates to and reproduces in the non-target population is 10^−3^, then the total number of male emigrants over the 36 generations is 0.013 (i.e., 1.3% of the original population), with a maximum value of 0.0005 in any one generation. The expected proportion of YLE males in the non-target population will then depend on its size relative to the target population. Assuming equal population sizes, the YLE is expected to be in 1.4% of males after 50 generations, and the number of females to be 98.3% of the number pre-release. If there is a small cost to the YLE (e.g., *s_Y_* = 0.05), and release rates increased to 5.3% to still achieve 95% suppression of the target population in 36 generations, then impacts on the non-target population are even lower, with a frequency in males after 50 generations of 0.4% and a female abundance of 99.5% (Fig. 6). If desired, increased fitness costs of a YLE could be engineered by adding extra sequences onto the Y or knocking-out some non-essential sequence. Tolerance thresholds for spread to neighboring populations will be determined on a case-by-case basis.

**Fig. 6.**
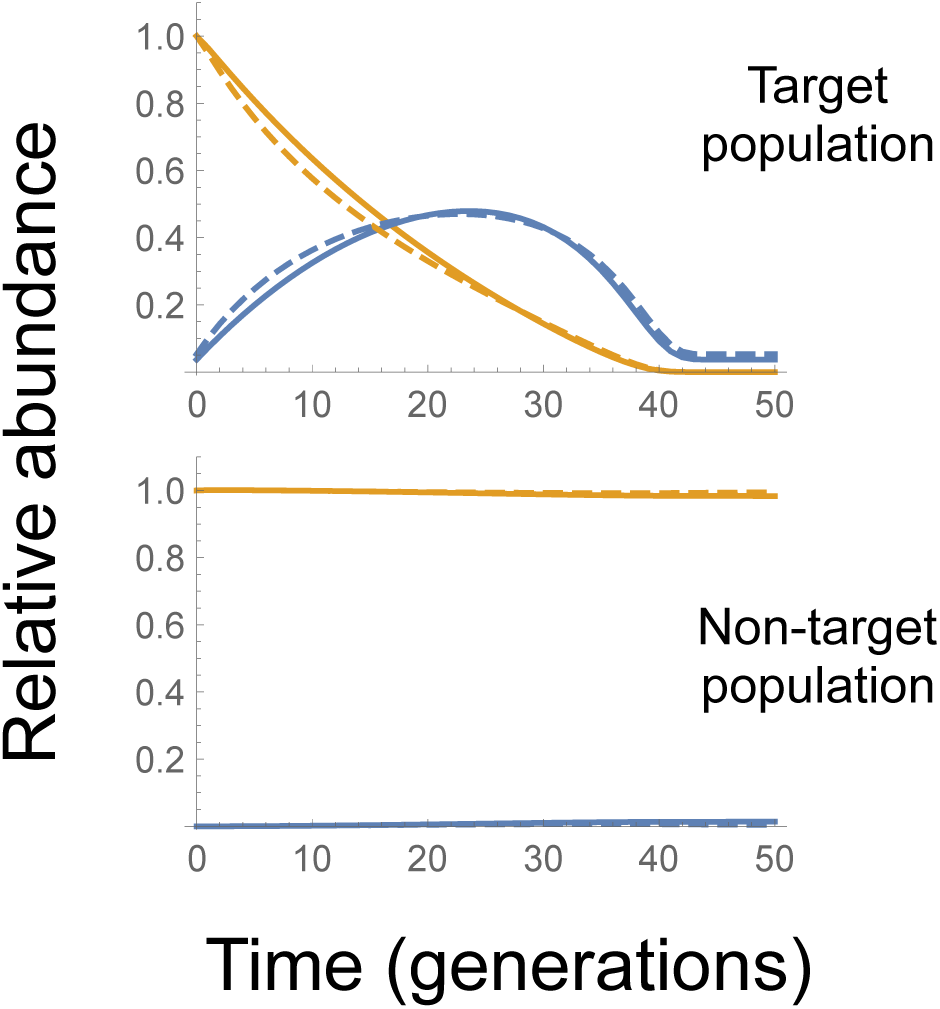
Frequency of YLE males (blue) and relative numbers of females (orange) in target and non-target populations over time. Solid lines show no cost of the YLE (*s_Y_*=0) with a release rate (that continues for the full duration of the simulation) of 3.7%, and dashed lines show *s_Y_*=0.05 with release rate of 5.3% — in both cases the release rates are sufficient to give 95% suppression in 36 generations. In every generation a pre-reproductive (and pre-mating) adult has a probability of 0.1% of moving from the target to non-target populations, regardless of gender or genotype. For simplicity, migration in the opposite direction is ignored.

#### Augmenting transmission of the YLE

All else being equal, the higher the frequency of the YLE in a population, the greater the load it imposes upon it. Is there any way to increase this frequency without releasing more males? In principle a YLE could be put on a driving Y chromosome, as modeled by Deredec et al. (Deredec et al. 2011), but then it may spread to and substantially suppress geographically distinct populations if there is even infrequent gene flow. Here we consider the possibility of releasing a YLE along with an autosomal X-shredder or other gene that leads to disproportionate transmission of the Y chromosome (Galizi et al. 2014, Galizi et al. 2016). Four possible implementations are considered, including an X-shredder that is either constitutive or conditional (i.e., causes biased transmission of all Ys or just of YLE-bearing Ys), and that is released either in the same males as the YLE, or in different males released at the same time. In each case the YLE targets an X-linked locus, and we initially model the impact of a single one-time release of 10% of the pre-release population.

Exemplar results for the four idealized scenarios are shown in Figure 7, along with no X-shredder release for comparison. With a constitutive X-shredder, if it is released in the same males as the YLE, it boosts the transmission of the YLE from one generation to the next, which therefore continues to increase in frequency even after releases have stopped (Fig. 7a, blue line), and as a result the population size continues to fall (black dashed line). The boost occurs because of the positive association, or linkage disequilibrium, between the X-shredder and the YLE (green line), which starts complete and then gradually decays over time, so the rate of increase of the YLE also slows. By contrast, if there are separate releases of YLE males and X-shredder males, then there is initially a negative correlation between the two loci, the X-shredder boosts the transmission of wildtype Ys, the frequency of the YLE declines over time, and the final level of control is worse than if no X-shredders were released (Fig. 7b vs. Fig. 7e,f).

**Fig. 7.**
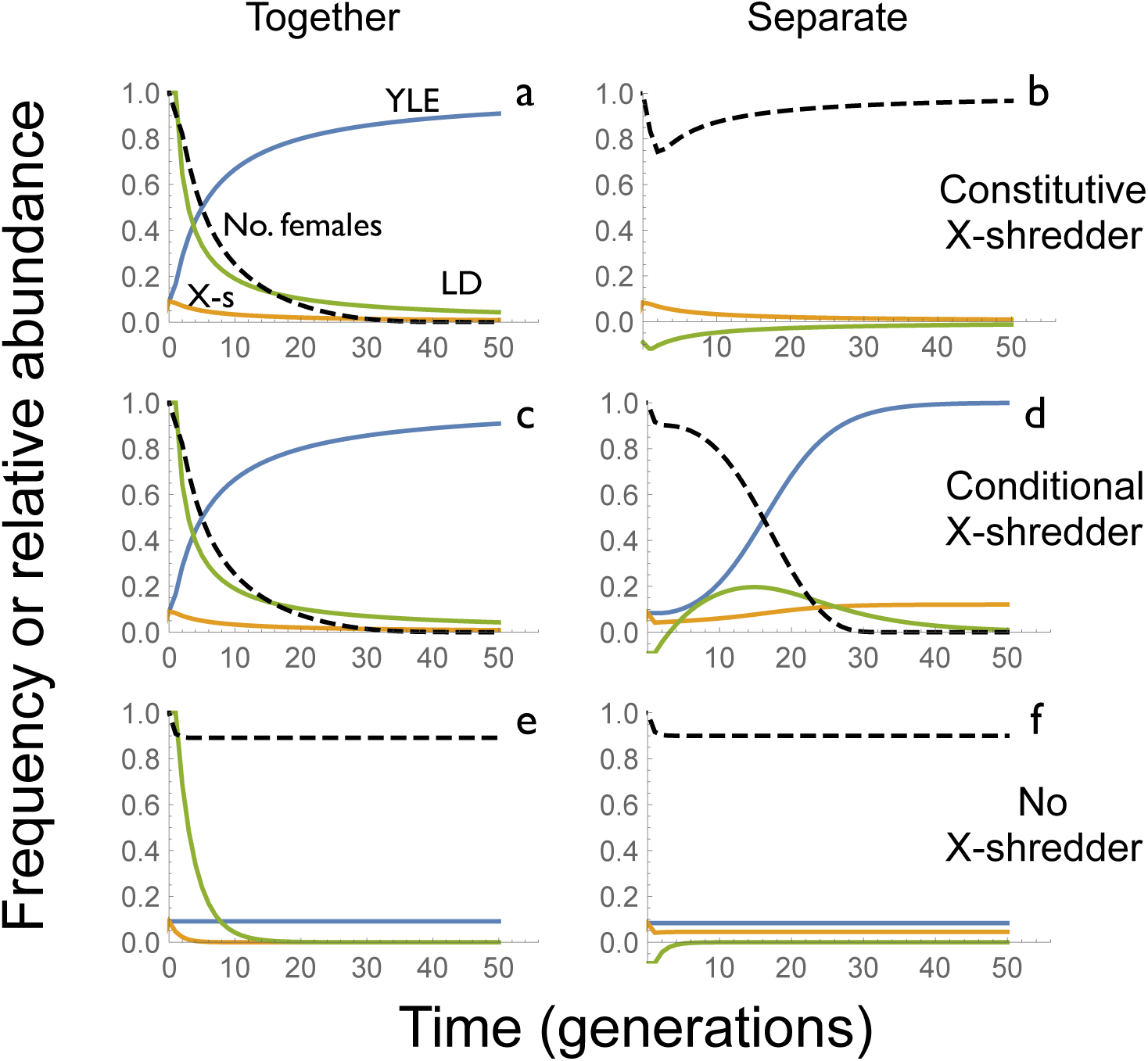
Impact of including an autosomal X-shredder in the releases. Scenarios considered include the X-shredder being present in the same released males as the YLE (left column), or in additional males that carry a wild-type Y (right column), and an X-shredder that causes 100% transmission of the Y regardless of the genotype of the Y (top row), or only of those carrying the YLE (middle row). Bottom row shows dynamics for a non-functional X-shredder (no impact on Y transmission). In all cases there is a single 10% release in generation 0 of males carrying the YLE that targets an X-linked locus; in the ‘separate’ scenarios there is an additional 10% release of males carrying the X-shredder. Black dashed lines shows abundance of females over time; blue line the frequency of the YLE in males; orange line the frequency of the X-shredder in males, and green line the correlation coefficient among males between the presence of the YLE and of the X-shredder (heterozygotes and homozygotes combined). All parameters at idealised values.

If instead the X-shredder only acts in the presence of the YLE, then it can boost the frequency of the YLE and the impact of a release whether it is initially in the same or different males, though the dynamics are different in detail (Fig. 7c,d). If they are released in the same males, then with the idealized parameter values considered here there is no difference in the dynamics whether the X-shredder is constitutive or conditional, because a 100% effective X-shredder is always transmitted to males, and remains associated with the Y chromosome with which it was released. Differences between constitutive and conditional X-shredders would appear with less than perfect performance. By comparison, if the YLE and X-shredder are released in separate males, then, at least in this example, the level of suppression is initially lower, but later it is higher. Importantly, in all these scenarios, though the YLE can substantially increase in frequency after release, the X-shredder shows either a decline or only a modest increase, indicating it may be unlikely to spread much into other geographically distinct populations, though a detailed analysis is beyond the scope of this study.

As the inclusion of an X-shredder causes the frequency of the YLE and the extent of suppression to continue to increase after a release, it may be more efficient to release once in a population rather than repeatedly. If we now ask how large a single release must be to achieve 95% suppression in 36 generations, it turns out to be 6.0% if the YLE-a and X-shredder are released in the same males, and 5.4% if the X-shredder is conditional and is released separately (i.e., 5.4% of YLE-a males and 5.4% of X-shredder males, assuming equal releases of the two types). In either case the total number of males that need to be released is more than an order of magnitude lower than without an X-shredder (3.6% releases for 36 generations; Table 1).

### Discussion

Our modeling has shown that YLEs can give substantially more efficient control of populations than the sterile insect technique or many other proposed variants, while still allowing the impact to be focused on a particular geographical region. This efficiency derives from the fact that YLEs are not selected against as a consequence of the harm they cause – they are evolutionarily insulated from their impacts (Deredec et al. 2011). YLEs share this feature with the “Trojan female” proposal of releasing into populations mitochondria that specifically impair male fertility (Wolff et al. 2017). Some naturally occurring maternally inherited endosymbionts in insects have even evolved to kill male embryos, to free up resources for their sisters (Hurst and Frost 2015), which highlights an important assumption of our models, that the YLE does not confer a fitness advantage to the male. Such an advantage could occur if brothers and sisters compete more intensely than random and the YLE caused early (e.g., embryonic) death of females. In such a situation the YLE could show a form of drive (Friberg and Rice 2015) and spread in a self-sustaining way, including to neighboring populations.

Other types of sex-linked genome editors are also possible and may be useful. We have considered the most obvious classes of target gene (lethals and steriles), but in some species there may be other possibilities, such as edits that convert females into males. In species where males play a more important role in provisioning for the next generation, or otherwise are more harmful, then it may be worth considering an X-linked editor that targets a Y-linked gene, including a male-determining gene. And in female heterogametic species, where females are WZ and males ZZ, if it is still desirable to release only males, then a Z-linked editor that targets a W-linked locus (including a female-determining gene) could be considered.

Our primary metric for the efficiency of an intervention strategy has been the release rates required to achieve a particular level of control. We have assumed the released males are equally fit as the wild males, though differences could arise because the released males are of lower vigor (e.g., due to poorer rearing environment, being inbred, or damage in transport), or they may not be released into the right locations at the right time. Such effects can be easily incorporated into our models, at least in a simple way: if released males have, say, one tenth the reproductive success of wild males, then the release rates calculated here need to be multiplied by 10. Incorporating low vigor due to lab adaptation may be somewhat more complex, as those traits would persist with the YLE for some generations. Most of our calculations have also assumed equal numbers released in each generation, but this may not be optimal. For example, it is possible that transportation costs are such that it will be better to release every second generation, or some other pattern. The fact that YLEs and their effects persist in the population better than other self-limiting constructs (Fig. 4) may allow greater operational flexibility. And as with many genetic strategies, impacts are determined by proportions, and, all else being equal, releasing into a small population will be more effective than releasing the same absolute number into a larger one. If the target population reproduces throughout the year, but shows seasonal cycles of abundance, then it may be most efficient to focus releases in the low season.

### Molecular possibilities

It may be possible to develop synthetic constructs with properties much like those modeled here in many different ways. First, one might insert onto the Y chromosome an editor that knocks out a gene that is haplo-insufficient for viability or fertility and is X-linked or has female-limited effects. If the target gene is autosomal and affects both sexes, then a “rescue” copy of the gene might be inserted on the Y that is sufficiently diverged in sequence that it is not recognized by the editor and, if the editor is a nuclease, could not act as a template for homologous repair. Knock-outs can be produced by sequence-specific cleavage followed by end-joining, in which case it may be useful to target sequences between direct repeats where micro-homology-mediated end joining would produce a frameshift mutation (Hammond et al. 2017). Conceivably one could insert a non-functional copy of the gene on the Y, to allow homologous repair, but it is not clear if this would be useful. Alternatively, instead of using a nuclease, one could use a base editor to, for example, change a C to a T and thereby introduce a premature stop codon (Billon et al. 2017, Komor et al. 2017).

Second, the YLE might target a gene that is haplo-sufficient for survival or fertility if, in addition to acting in the male germline, there is also paternal carry-over of the editing complex via the sperm into the zygote, where it then disrupts the maternally derived allele. Paternal carry-over of the PpoI meganuclease has been observed in the mosquito *Anopheles gambiae* (Windbichler et al. 2008, Galizi et al. 2014). Again, the target gene would need to have female-limited effects, or there could be a rescue copy on the Y, and the knock out could be produced by a nuclease or a base editor. In principle it may also be possible to introduce dominant negative mutations into a haplo-sufficient gene, in which case the paternal carry-over would not be necessary.

Third, the YLE might target the X chromosome in a number of other ways. It might cleave a sequence on the X, followed by homologous repair from the Y that inserts a female-specific dominant lethal gene. In some species inserting a copy of the male determining gene may be sufficient (Criscione et al. 2016, Krzywinska et al. 2016). This approach requires some homology between X and Y, which could pre-exist (e.g., in a pseudo-autosomal region), or could be engineered. Or, the YLE might encode a protein that binds to the X and is carried with it into the zygote, where it causes death – a poison tag system that may, for example, use chromatin regulators (Keung et al. 2015, Park et al. 2016).

Regardless of what sequence is targeted by the YLE, it will be important to consider the potential for target site resistance to evolve (Beaghton et al. 2017a, Champer et al. 2017, Hammond et al. 2017), and control sequences will need to be chosen to ensure the YLE is active in the male germline. If the edit has female-limited effects then some somatic expression may be acceptable, but if the target is on the X and there is no rescue copy on the Y then somatic expression may be harmful to the male and should be minimized. Similarly, expression may need to be as late as possible in the male germline if the target gene is expressed there. In some species the sex chromosomes are silenced around the time of meiosis, in which case expression may need to occur earlier (e.g., (Bernardini et al. 2014)) or the silencing circumvented. Rearing the organisms for release is likely to be easier if the YLE can be made repressible in the production facility (Thomas et al. 2000).

Other alternatives exist outside the YLE paradigm. For example, a Y-linked spermatogenically expressed toxin and zygotically expressed antidote, analogous to those developed for synthetic underdominance and medea systems (Akbari et al. 2013, Akbari et al. 2014), could be used. A paternally active toxin-antidote system has been discovered in natural populations of the nematode *Caenorhabditis elegans* (Seidel et al. 2011). And, our modeling suggests that an autosomal construct showing dominant female lethality and drive in males (Thomas et al. 2000, Adelman and Tu 2016, Piaggio et al. 2017) can be as efficient as a YLE. In the future it would be useful to compare these different molecular alternatives in terms of other criteria, including evolutionary stability in the face of mutations likely to arise after release (Burt 2003, Beaghton et al. 2017b).

Combining a YLE with an autosomal X-shredder, or other gene that distorts transmission of the sex chromosomes, can be even more efficient. Both meganuclease- and CRISPR-based X-shredders that act by targeting the rDNA repeat have been developed in *An. gambiae* (Galizi et al. 2014, Galizi et al. 2016). A conditional CRISPR-based X-shredder that only acts in the presence of the YLE could be designed by separating the Cas9 and guide RNA genes, inserting one on the Y and the other on an autosome. The efficiency of these “augmented YLEs” could presumably be further enhanced by adding a third locus that increases the transmission of the X-shredder, as has been shown with the “daisy drives” of Noble et al. (Noble et al. 2016), though at the cost of increased genetic (and regulatory) complexity. Increased efficiency may also be achieved by placing the X-shredder in a pseudo-autosomal region (if such exists or could be constructed), where linkage with the YLE could be tighter than if it is on an autosome, though some mechanism may then be needed to prevent an inversion creating a driving Y.

Recent advances in molecular biology and genome editing are opening up new possibilities for the safe and effective control of harmful species. The species for which genetic approaches are being considered vary widely, from disease-transmitting mosquitoes to invasive mammals, and it is unlikely that the same approach will be appropriate in all cases. YLEs offer a unique combination of features that may be useful in some applications, particularly if efficient control is needed in some locations and is to be avoided in others.Whether YLEs are useful in any particular application, and the design criteria needed for success, will require more tactical models tailored to the biology of the target species.

## The Model

### Genetics and fitness effects

The multiple diverse strategies are investigated with three genetic models. In the first, there are two Y chromosomes, wild-type and transgenic, denoted y and Y, and an autosomal locus with two alleles, wildtype and variant (mutant or transgenic, depending on the scenario), denoted a and A. There are thus 3 female genotypes (a/a, a/A, and A/A), producing 2 types of egg (a and A), and 6 male genotypes (ya/a, ya/A, yA/A, Ya/a, Ya/A, and YA/A), producing 4 types of sperm (ya, yA, Ya, YA). In a/A heterozygotes, the A allele is transmitted to a fraction *df* of eggs and *dm* of sperm (*df*, *dm* = 0.5 for Mendelian transmission). In a/A males the Y chromosome is transmitted to a fraction *m1* of progeny, and in A/A males to a fraction *m2* of progeny (*m1*=*m2*=0.5 for Mendelian transmission). In males with a transgenic Y chromosome and at least one a allele at the autosomal locus, the a allele mutates to A with probability *u*; these mutations are assumed to occur in the germline of the male and have no effect on his survival or fertilization success. Table SI-1 shows the parameter settings for the various strategies, and Table SI-2 shows the proportions of gametes produced by the different genotypes.

In the second model, used for a YLE targeting an X-linked locus, the second locus is on the X chromosome instead of an autosome, with two alleles, wildtype and variant, denoted x and X. There are 3 female genotypes (x/x, x/X, and X/X), producing 2 types of egg (x and X), and 4 male genotypes (yx, yX, Yx and YX), producing 4 types of sperm (x, X, y and Y). Transmission is Mendelian except for the mutations of x to X in the presence of Y (Table SI-3). To incorporate an X-shredder, a third (autosomal) locus is added to the model, which determines the transmission rates of the sex chromosomes.

### Population biology and selection

We model a population with discrete, non-overlapping generations. In the pre-intervention wildtype population each generation begins with a certain number of hatchlings, denoted N_h_. Non-selective density-dependent mortality occurs during the juvenile phase, such that the probability of surviving is equal to θ_J_ α/ (α + N_h_[t]), where θ_J_ is the density-independent (or low density) probability a hatchling survives to become an adult; *α* is a constant determining the intensity of density-dependent mortality; and N_h_[t] is the number of hatchlings (male plus female) at time t. Individuals then become adults and mate randomly, with each wildtype female producing *f* fertilised eggs to start the next generation. Each juvenile is derived from an independent mating event, and males are assumed not to be limiting in the production of fertilized eggs. The intrinsic rate of increase of the population is

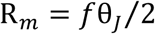

and the pre-intervention equilibrium population size is

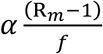

Selection on the autosomal or X-linked locus (i.e., differential survival or reproduction by genotype) can occur in one of three ways: (i) differential survival of hatchlings before the density-dependent mortality; (ii) differential survival of pre-adults after the density-dependent mortality; or (iii) differential fecundity of adult females and mating success of adult males. Whichever of these occurs, the fitness of wildtype genotypes (a/a or x/x) is standardised to 1; the fitness of heterozygotes (a/A or x/X) is 1 − h_f_ s_f_ for females and 1 − h_m_ s_m_ for males; and the fitness of homozygotes (A/A or X/X) or hemizygotes (X males) is 1 − s_f_ for females and 1 − s_m_ for males, where s_f_ and s_m_ are selection coefficients and h_f_ and h_m_ are dominance coefficients. In addition, males with a transgenic Y have fitness reduced by a factor 1 − s_Y_; for simplicity this selection assumed to occur at the same time and in the same manner as selection on the autosomal or X-linked locus. Idealized parameter values for the various selection and dominance coefficients are shown in Table SI-1, and fitnesses of the different genotypes shown in Tables SI-2 and SI-3.

---

**Table SI-1.** Idealised parameter settings for different genetic control strategies.

| Strategy | Transmission rates | | | | Mutation<br>rate ($u$) | Fitness parameters | | | | | Genotype<br>released |
| --- | --- | --- | --- | --- | --- | --- | --- | --- | --- | --- | --- |
| | $d_m$ | $d_f$ | $m_1$ | $m_2$ | | $h_f$ | $s_f$ | $h_m$ | $s_m$ | $s_Y$ | |
| bi-RIDL | 1/2 | 1/2 | 1/2 | 1/2 | 0 | 1 | 1 | 1 | 1 | - | yA/A |
| fs-RIDL | 1/2 | 1/2 | 1/2 | 1/2 | 0 | 1 | 1 | 0 | 0 | - | yA/A |
| fs-RIDL-drive | 1 | 1 | 1/2 | 1/2 | 0 | 1 | 1 | 0 | 0 | - | ya/A |
| X-shredder | 1/2 | 1/2 | 1 | 1 | 0 | 0 | 0 | 0 | 0 | - | yA/A |
| Y-linked editor | 1/2 | 1/2 | 1/2 | 1/2 | 1 | 1 | 1 | 0 | 0 | 0 | Ya/a |

$d_m$ and $d_f$ are transmission rates of the A allele in a/A heterozygous males and females, respectively. $m_1$ and $m_2$ are transmission rates of the Y chromosome (whether transgenic or not) in a/A and A/A males, respectively. $u$ is the rate at which a alleles mutate to A in transgenic Y males. $s_f$ and $s_m$ are selection coefficients against homozygous A/A females and males, $h_f$ and $h_m$ are the corresponding dominance coefficients, and $s_Y$ is the selection coefficient against the Y-linked editor gene.

**Table SI-2.** The fitness of each genotype and the proportion of each gamete produced by them for the model of polymorphic Y chromosome and autosome

| Diploid<br>genotype | Fitness | Gametes produced |  |  |  |  |  |
| --- | --- | --- | --- | --- | --- | --- | --- |
| | | $xa$ | $xA$ | $ya$ | $yA$ | $Ya$ | $YA$ |
| $xaa$ | 1 | 1 | 0 | 0 | 0 | 0 | 0 |
| $xaA$ | $1-h_f s_f$ | $1-d_f$ | $d_f$ | 0 | 0 | 0 | 0 |
| $xAA$ | $1-s_f$ | 0 | 1 | 0 | 0 | 0 | 0 |
| $yaa$ | 1 | 1/2 | 0 | 1/2 | 0 | 0 | 0 |
| $yaA$ | $1-h_m s_m$ | $(1-d_m)(1-m_1)$ | $d_m(1-m_1)$ | $(1-d_m)m_1$ | $d_m m_1$ | 0 | 0 |
| $yAA$ | $1-s_m$ | 0 | $1-m_2$ | 0 | $m_2$ | 0 | 0 |
| $Yaa$ | $1-s_Y$ | $(1-u)/2$ | $u/2$ | 0 | 0 | $(1-u)/2$ | $u/2$ |
| $YaA$ | $(1-s_Y)(1-h_m s_m)$ | $(1-u)/4$ | $(1+u)/4$ | 0 | 0 | $(1-u)/4$ | $(1+u)/4$ |
| $YAA$ | $(1-s_Y)(1-s_m)$ | 0 | 1/2 | 0 | 0 | 0 | 1/2 |

**Table SI-3.** The fitness of each genotype and the proportion of each gamete produced by them for the model of polymorphic Y and X chromosomes

| Diploid<br>genotype | Fitness | Gametes produced |  |  |  |
| --- | --- | --- | --- | --- | --- |
| | | $x$ | $X$ | $y$ | $Y$ |
| $xx$ | 1 | 1 | 0 | 0 | 0 |
| $xX$ | $1-h_f s_f$ | 1/2 | 1/2 | 0 | 0 |
| $XX$ | $1-s_f$ | 0 | 1 | 0 | 0 |
| $yx$ | 1 | 1/2 | 0 | 1/2 | 0 |
| $yX$ | $1-s_m$ | 0 | 1/2 | 1/2 | 0 |
| $Yx$ | $1-s_Y$ | $(1-u)/2$ | $u/2$ | 0 | 1/2 |
| $YX$ | $(1-s_Y)(1-s_m)$ | 0 | 1/2 | 0 | 1/2 |

## Literature cited

Adelman, Z. N. and Z. J. Tu (2016). Control of mosquito-borne infectious diseases: sex and gene drive. Trends Parasitol. 32(3): 219–229.

Akbari, O. S., C.-H. Chen, J. M. Marshall, H. Huang, I. Antoshechkin and B. A. Hay (2014). Novel synthetic medea selfish genetic elements drive population replacement in *Drosophila;* a theoretical exploration of medea-dependent population suppression. Acs Synthetic Biology 3(12): 915–928.

Akbari, O. S., K. D. Matzen, J. M. Marshall, H. Huang, C. M. Ward and B. A. Hay (2013). A synthetic gene drive system for local, reversible modification and suppression of insect populations. Current Biology 23(8): 671–677.

Alphey, L. (2014). Genetic control of mosquitoes. Ann. Rev. Entomol. 59: 205–224.

Alphey, L., M. Benedict, R. Bellini, G. G. Clark, D. A. Dame, M. W. Service and S. L. Dobson (2010). Sterile-insect methods for control of mosquito-borne diseases: an analysis. Vector-Borne Zoonotic Dis. 10(3): 295–311.

Alphey, N. and M. B. Bonsall (2014). Interplay of population genetics and dynamics in the genetic control of mosquitoes. Journal of the Royal Society Interface 11(93).

Beaghton, A., P. J. Beaghton and A. Burt (2016). Gene drive through a landscape: reaction-diffusion models of population suppression and elimination by a sex ratio distorter. Theoretical Population Biology 108: 51–69.

Beaghton, A., P. J. Beaghton and A. Burt (2017a). Vector control with driving Y chromosomes: modelling the evolution of resistance. Malaria Journal 16.

Beaghton, A., A. Hammond, T. Nolan, A. Crisanti, H. C. J. Godfray and A. Burt (2017b). Requirements for driving antipathogen effector genes into populations of disease vectors by homing. Genetics 205(4): 1587–1596.

Bernardini, F., R. Galizi, M. Menichelli, P. A. Papathanos, V. Dritsou, E. Marois, A. Crisanti and N. Windbichler (2014). Site-specific genetic engineering of the *Anopheles gambiae* Y chromosome. Proc. Natl. Acad. Sci. USA 111(21): 7600–7605.

Billon, P., E. E. Bryant, S. A. Joseph, T. S. Nambiar, S. B. Hayward, R. Rothstein and A. Ciccia (2017). CRISPR-mediated base editing enables efficient disruption of eukaryotic genes through induction of STOP codons. Mol. Cell 67(6): 1068-+.

Burt, A. (2003). Site-specific selfish genes as tools for the control and genetic engineering of natural populations. Proc. Roy. Soc. Lond. B 270(1518): 921–928.

Burt, A. (2014). Heritable strategies for controlling insect vectors of disease. Philosophical Transactions of the Royal Society B-Biological Sciences 369(1645).

Carvalho, D. O., A. R. McKemey, L. Garziera, R. Lacroix, C. A. Donnelly, L. Alphey, A. Malavasi and M. L. Capurro (2015). Suppression of a field population of *Aedes aegypti* in Brazil by sustained release of transgenic male mosquitoes. PLoS Neglected Tropical Diseases 9(7).

Champer, J., R. Reeves, S. Y. Oh, C. Liu, J. X. Liu, A. G. Clark and P. W. Messer (2017). Novel CRISPR/Cas9 gene drive constructs reveal insights into mechanisms of resistance allele formation and drive efficiency in genetically diverse populations. Plos Genetics 13(7).

Criscione, F., Y. M. Qi and Z. J. Tu (2016). GUY1 confers complete female lethality and is a strong candidate for a male-determining factor in *Anopheles stephensi*. Elife 5.

Deredec, A., H. C. J. Godfray and A. Burt (2011). Requirements for effective malaria control with homing endonuclease genes. Proc. Natl. Acad. Sci. USA 108(43): E874–E880.

Dhole, S., M. R. Vella, A. L. Lloyd and F. Gould (2017). Invasion and migration of spatially self-limiting gene drives: a comparative analysis. Biorxiv.

Eckhoff, P. A., E. A. Wenger, H. C. J. Godfray and A. Burt (2017). Impact of mosquito gene drive on malaria elimination in a computational model with explicit spatial and temporal dynamics. Proc. Natl. Acad. Sci. USA 114(2): E255–E264.

Friberg, U. and W. R. Rice (2015). Sexually antagonistic zygotic drive: a new form of genetic conflict between the sex chromosomes. Cold Spring Harbor Perspectives in Biology 7(3).

Fu, G., R. S. Lees, D. Nimmo, D. Aw, L. Jin, P. Gray, T. U. Berendonk, H. White-Cooper, S. Scaife, H. K. Phuc, O. Marinotti, N. Jasinskiene, A. A. James and L. Alphey (2010). Female-specific flightless phenotype for mosquito control. Proc. Nat. Acad. Sci. USA 107: 4550–4554.

Galizi, R., L. A. Doyle, M. Menichelli, F. Bernardini, A. Deredec, A. Burt, B. L. Stoddard, N. Windbichler and A. Crisanti (2014). A synthetic sex ratio distortion system for the control of the human malaria mosquito. Nature Communications 5.

Galizi, R., A. Hammond, K. Kyrou, C. Taxiarchi, F. Bernardini, S. M. O’Loughlin, P. A. Papathanos, T. Nolan, N. Windbichler and A. Crisanti (2016). A CRISPR-Cas9 sex-ratio distortion system for genetic control. Scientific Reports 6.

Godfray, H. C. J., A. North and A. Burt (2017). How driving endonuclease genes can be used to combat pests and disease vectors. BMC Biology 15.

Hammond, A., R. Galizi, K. Kyrou, A. Simoni, C. Siniscalchi, D. Katsanos, M. Gribble, D. Baker, E. Marois, S. Russell, A. Burt, N. Windbichler, A. Crisanti and T. Nolan (2016). A CRISPR-Cas9 gene drive system-targeting female reproduction in the malaria mosquito vector *Anopheles gambiae*. Nature Biotechnology 34(1): 78–83.

Hammond, A. M., K. Kyrou, M. Bruttini, A. North, R. Galizi, X. Karlsson, N. Kranjc, F. M. Carpi, R. D’Aurizio, A. Crisanti and T. Nolan (2017). The creation and selection of mutations resistant to a gene drive over multiple generations in the malaria mosquito. PLoS Genetics 13(10).

Hurst, G. D. D. and C. L. Frost (2015). Reproductive parasitism: maternally inherited symbionts in a biparental world. Cold Spring Harbor Perspectives in Biology 7(5).

Keung, A. J., J. K. Joung, A. S. Khalil and J. J. Collins (2015). Chromatin regulation at the frontier of synthetic biology. Nature Reviews Genetics 16(3): 159–171.

Komor, A. C., K. T. Zhao, M. S. Packer, N. M. Gaudelli, A. L. Waterbury, L. W. Koblan, Y. B. Kim, A. H. Badran and D. R. Liu (2017). Improved base excision repair inhibition and bacteriophage Mu Gam protein yields C:G-to-T: A base editors with higher efficiency and product purity. Science Advances 3(8).

Krzywinska, E., N. J. Dennison, G. J. Lycett and J. Krzywinski (2016). A maleness gene in the malaria mosquito *Anopheles gambiae*. Science 353(6294): 67–69.

Mains, J. W., C. L. Brelsfoard, R. I. Rose and S. L. Dobson (2016). Female adult *Aedes albopictus* suppression by *Wolbachia-infected* male mosquitoes. Scientific Reports 6.

Marshall, J. M. and B. A. Hay (2012). Confinement of gene drive systems to local populations: a comparative analysis. J. Theor. Biol. 294: 153–171.

Noble, C., J. Min, J. Olejarz, J. Buchthal, A. Chavez, A. L. Smidler, E. A. DeBenedictis, G. M. Church, M. A. Nowak and K. M. Esvelt (2016). Daisy-chain gene drives for the alteration of local populations. bioRxiv. http://dx.doi.org/10.1101/057307.

Park, M., A. J. Keung and A. S. Khalil (2016). The epigenome: the next substrate for engineering. Genome Biology 17.

Phuc, H., M. H. Andreasen, R. S. Burton, C. Vass, M. J. Epton, G. Pape, G. Fu, K. C. Condon, S. Scaife, C. A. Donnelly, P. G. Coleman, H. White-Cooper and L. Alphey (2007). Late-acting dominant lethal genetic systems and mosquito control. BMC Biology 5(11): (20 March 2007).

Piaggio, A. J., G. Segelbacher, P. J. Seddon, L. Alphey, E. L. Bennett, R. H. Carlson, R. M. Friedman, D. Kanavy, R. Phelan, K. H. Redford, M. Rosales, L. Slobodian and K. Wheeler (2017). Is it time for synthetic biodiversity conservation? Trends Ecol. Evol. 32(2): 97–107.

Prout, T. (1978). Joint effects of release of sterile males and immigration of fertilized females on a density regulated population. Theor. Pop. Biol. 13(1): 40–71.

Schliekelman, P., S. Ellner and F. Gould (2005). Pest control by genetic manipulation of sex ratio. J. Econ. Entomol. 98: 18–34.

Schliekelman, P. and F. Gould (2000). Pest control by the release of insects carrying a female-killing allele on multiple loci. J. Econ. Entomol. 93(6): 1566–1579.

Seidel, H. S., M. Ailion, J. L. Li, A. van Oudenaarden, M. V. Rockman and L. Kruglyak (2011). A novel sperm-delivered toxin causes late-stage embryo lethality and transmission ratio distortion in *C. elegans*. PLoS Biology 9(7).

Thomas, D. D., C. A. Donnelly, R. J. Wood and L. S. Alphey (2000). Insect population control using a dominant, repressible, lethal genetic system. Science 287(5462): 2474–2476.

Windbichler, N., P. A. Papathanos and A. Crisanti (2008). Targeting the X chromosome during spermatogenesis induces Y chromosome transmission ratio distortion and early dominant embryo lethality in *Anopheles gambiae*. PLoS Genet. 4(12): e1000291, 1000291–1000299.

Wolff, J. N., N. J. Gemmell, D. M. Tompkins and D. K. Dowling (2017). Introduction of a male-harming mitochondrial haplotype via ‘Trojan Females’ achieves population suppression in fruit flies. Elife 6.

Yakob, L., L. Alphey and M. B. Bonsall (2008). *Aedes aegypti* control: the concomitant role of competition, space and transgenic technologies. J. Appl. Ecol. 45(4): 1258–1265.

Zhang, D. J., R. S. Lees, Z. Y. Xi, K. Bourtzis and J. R. L. Gilles (2016). Combining the sterile insect technique with the incompatible insect technique: III-Robust mating competitiveness of irradiated triple *Wolbachia-infected Aedes albopictus* males under semi-field conditions. PLoS One 11(3).

